# Recombinant expression of osmotin in barley improves stress resistance and food safety during adverse growing conditions

**DOI:** 10.1101/546721

**Authors:** Jitka Viktorova, Barbora Klcova, Katerina Rehorova, Tomas Vlcko, Lucie Stankova, Nikola Jelenova, Pavel Cejnar, Jiban Kumar Kundu, Ludmila Ohnoutkova, Tomas Macek

## Abstract

Although many genetic manipulations of crops providing biofortified or safer food have been prepared, the acceptance of biotechnology crops still remains limited. We report on a transgenic barley expressing the multi-functional protein osmotin that improves plant defense under stress conditions. An Agrobacterium–mediated technique was used to transform immature embryos of the spring barley cultivar Golden Promise. Transgenic barley plants of the T0 and T1 generations were evaluated by molecular methods.

Transgenic barley tolerance to stress was determined by chlorophyll, total protein, malondialdehyde and ascorbate peroxidase content. Transgenic plants maintained the same level of chlorophyll and protein, which significantly declined in wild-type barley under the same stressful conditions. Salt stress evoked higher ascorbate peroxidase level and correspondingly less malondialdehyde. Methanol extracts of i) *Fusarium oxysporum* infected or ii) salt-stressed plants, were characterized by their acute toxicity effect on human dermal fibroblasts (HDF). Osmotin expressing barley extracts exhibited a lower cytotoxicity effect of statistical significance than that of wild-type plants under both types of stress tested on human dermal fibroblasts. Extract of *Fusarium oxysporum* infected transgenic barley was not able to damage DNA in Comet assay, which is in opposite to control plants. Moreover, this particular barley did not affect the local biodiversity interactions, which was tested through monitoring barley natural virus pathogen – host interactions – the BYDV and WDV viruses transmitted to the plants by aphids and leafhoppers. Our findings provide a new perspective which could help to evaluate the safety of products from genetically modified crops.

## 1 Introduction

Osmotin is a 26-kDa protein belonging to the 5^th^ class of pathogenesis-related proteins (PR 5) together with thaumatin, zeamatin and others. Osmotin was first characterized in *Nicotiana tabacum* plants adapted to saline conditions [1, 2]. Later, its homologs were found in both monocots and dicots across the whole plant kingdom [3]. Osmotin is a multifunctional protein that plays an important role in the plant immune system during stress. Abiotic originators of stress such as drought, salt and cold induce osmotin expression. As the name indicates, osmotin plays a role as an osmoprotectant, providing enzyme protection and protein chaperone functions. Biotic stress resistance is related to the ability of osmotin to activate the fungal receptor that induces the programmed cell death of fungi. Being a promising protein, it has served as a tool for biotechnology engineering for many decades [1].

The first transgenic plants containing the heterologous osmotin gene in their genomes were potato and tobacco [4] and many others followed, as summarized in reviews [3, 5]. More recently, transgenic monocotyledonous plants bearing the osmotin gene were prepared as well. Osmotin was first expressed by a monocot wheat in 2008 [6] followed by rice another three years later [7]. The genetic manipulation of monocots is a promising and still developing field, because their agriculture is significantly endangered by climatic changes. Monocots play an important role in agriculture for animal feed and for human nutrition, as they are among the most often planted crops (*e.g.* wheat, barley, rice, maize and others), however, the application of their GM variants still remains limited.

Even though transgenic crops are one of the most characterized and toxicologically tested plants, many of them have never been applied (*e.g*. Golden Rice [8]). The economic aspects, environmental benefits and higher yield of GM crops [9] usually do not affect public opinion significantly. Even though biofortified crops (reviewed *e.g*. in [10]) with improved properties and safer food (*e.g*. [11]) were reported years ago, the usage of GM crops still remains limited. No conventionally bred crop has been so thoroughly tested for toxicity, allergenicity or effect on the environment, intestinal microbiome and nutrition. However, what is often neglected is the increased toxicity of crops protecting themselves against harsh environmental conditions.

As a protection against stress, plants have adapted their own immune system [12] in order to survive. However, many of the secondary metabolites formed this way can be toxic compounds, *e.g*. glycoalkaloids [13], steroidal alkaloids [14], flavonoids [15, 16] and glycosides [17, 18]. Moreover, the toxicity of plant immune system compounds is often supplemented by the toxicity of the pathogen’s secondary metabolites, *e.g*. mycotoxins [19].

The aim of this work is the preparation of a transgenic barley with improved resistance to stress originators, documentation of this higher resistance by biochemical methods and proof of better toxicological properties for consumers compared to commercially used crops. As an often-mentioned disadvantage of GM plants, its influence on biodiversity is described. Here, we also focused on studying the effect of this GM barley on the spreading of viruses by aphids and leafhoppers. Two economically most important virus pathogens for barley cereals in Europe were selected – Barley yellow dwarf virus, transmitted by aphids, and Wheat dwarf virus, transmitted by leafhoppers.

## 2 Materials and methods

### 2.1 Barley transformation

The osmotin gene (*OSM*, GenBank: M29279.1, NCBI) was synthetized artificially (GeneArtTM Gene Synthesis, Thermo Fisher Scientific), therefore codon optimization for barley was performed. The osmotin gene was further cloned into the vector pDONR207 (Invitrogen) by BP clonase reaction (Gateway®, Thermo Fisher Scientific). The construct was verified by restriction. The osmotin gene was inserted by LR clonase reaction (Gateway® LR ClonaseTM, Thermo Fisher Scientific) into the destination vector pBract214 (http://www.bract.org). The pBract214 vector was designed for the transformation of barley. The gene of interest is under the control of the maize Ubi promoter. The vector contains the *hpt* gene conferring hygromycin resistance under the CaMV 35S promoter. Correct orientation of the transgene was verified by restriction analysis and sequencing. The Vector pBract214::*osm* (**Supplementary Figure 1**) and the helper plasmid pSoup were transformed into the *Agrobacterium tumefaciens* strain AGL1 by electroporation. The *Agrobacterium*-mediated transformation of the immature zygotic barley embryo genotype ‘Golden Promise’ was performed according to the transformation protocol by Harwood et al [20]. Explants were cultivated *in vitro* on selection - callus induction, and regeneration medium and transferred into soil. Putative transgenic T0 plants were screened by PCR analysis. The analysis was performed with genomic DNA that was isolated from leaf tissue of the regenerated plants. For PCR reaction, premix REDTaq® ReadyMixTM PCR Reaction Mix (Sigma-Aldrich, USA,) was used. The presence of the osmotin gene was determined by amplifying a 222-bp fragment using the primers F: 5’-GCCCTGCCTTCATACGCTAT-3’ and R: 5’-TACGGGCAGTTGTTCCTCAC-3’. The presence of the *hpt* selection gene by amplifying a 275-bp amplicon using the primers F: 5’-GATTGCTGATCCCCATGTGT-3’and R: 5’-GCTGCTCCATACAAGCCAAC-3’.

Transgene expression was verified at the mRNA level. Where not stated otherwise, all procedures were done according to the manufacturer’s instructions. RNA was extracted from young leaf tissue of the transgenic plant with an RNAqueous Total RNA Isolation Kit (Thermo Fisher Scientific). The sample was treated with a Turbo DNA-free TM Kit (Thermo Fisher Scientific) and RNA concentration was assessed spectrophotometrically (DeNovix, DS-11 Spectrophotometer). 1 ug of total RNA was reverse transcribed using RevertAidTM H minus Reverse transcriptase (Thermo Fisher Scientific). To analyze the reaction efficiency, dilution series of the selected cDNA samples were prepared. The endogenous gene for elongation factor [21] was selected as an internal control. For mRNA expression verification, a SensiFASTTM SYBR® No-ROX Kit (Bioline) was used. Three-step PCR was conducted using a MyGo Mini real-time cycler (IT-IS Life Science Ltd.). Primer sequences for osmotin transcript detection were F: 5’-TCAGGTCCAGCTTCGTGTTC-3’and R: 5’-TACGGGCAGTTGTTCCTCAC-3’and produced an amplicon of 85 bp. Initial denaturation at 95 °C for 180 s was followed by forty cycles of denaturation at 95 °C, annealing at 60 °C and elongation at 72 °C. Reaction was terminated by a final 5 min extension at 72 °C. Melting analysis and electrophoretic separation of PCR products were done to verify primer specificity. Transgenic barley plants with verified expression of osmotin gene were grown in greenhouse to maturity. Immature embryos (T1 generation) were dissected from young caryopses of T0 plants and were selected on half-MS medium containing hygromycin 75 mg/L. Subsequently, the germinating plants were transferred to pots. All the germinated plants were analyzed for the presence of the osmotin and *hpt* transgenes by PCR.

### 2.2 Stressing of transgenic and control plants

Isolated embryos of the T1 generation of obtained GM plants and of the non-GM barley were sown in individual pots (10 x 10 cm) filled with universal gardening substrate and placed in growing chambers under controlled conditions (temperature 24/16°C, air humidity 40 %, 16/8 day/night). Until 50 days after sowing, all plants had four developed leaves and were divided into groups according to the planned stress treatment. Each group consisted of 4 transgenic and 4 non-transgenic plants. The control group was planted under the same conditions, another group was irrigated with 200 mM NaCl every second day, the last group was treated with spores of *Fusarium oxysporum* DBM 4199 (OD=0.7) once per week. One fully developed leaf was cut off every 3 days and kept at −80 °C until the biochemical analysis. 16 days after the beginning of the stress conditions, all biomass was harvested and lyophilized for toxicity testing.

The BYDV infection was implemented using the aphids *Rhopalosiphum padi* carrying the BYDV-PAV species [22]. At the stage of two unfolded leaves, approximately five viruliferous aphids were transmitted onto each of 6 transgenic and 6 control group plants. The aphids were maintained there for four days, after that, the plants were treated with insecticide (Mospilan, Nippon Soda Co.) and kept under controlled conditions (temperature 19/15°C, 16/8 day/night).

The WDV virus infection was administered using leafhoppers *Psammotettix alienus* carrying the WDV-B barley strain [22]. At the stage of the main shoot and one tiller being detectable, approximately five viruliferous leafhoppers were transmitted onto each of 6 transgenic and 6 control group plants. The leafhoppers were maintained there for fourteen days, after that the leafhoppers were removed by hand and the plants were closely checked for any missing leafhoppers. The plants were then kept under controlled conditions as before (temperature 19/15°C, 16/8 day/night).

Both WDV and BYDV isolates, as well as infected aphids and leafhoppers were obtained from the Virus Collection of the Crop Research Institute, Prague (Virus Collection, 2017). The aphid and the leafhopper count of insect vectors required for successful infection were estimated based on the longterm experience with testing of cereal cultivars in the Crop Reseach Institute, Prague [23, 24].

### 2.3 Biochemical assays

For chlorophyll content determination, 0.02 g of barley leaf tissue was used. Samples were incubated in 99 % alcohol (1:100 (w/V)) in the dark for 24 h. The subsequent measurement was the same as described in the method by [25]. The preparation of samples for the determination of protein content and enzyme activities was universal in order to decrease sample weight and consumption of plant material. 0.1 g of leaf tissue was homogenized in liquid nitrogen into powder. Then, 2 ml of extraction buffer (50 mM phosphate buffer, 1 mM EDTA and 2 % (w/v) polyvinylpyrrolidone of pH 7.8) was added. The homogenized sample was centrifuged at 14,000 × g at 4 °C for 30 min. Protein content was measured according to the method by Bradford [26]. The reaction mixture contained plant extract, extraction buffer and reagent in the ratio 1:1:8. The reagent and subsequent measurement was the same as in the original paper. The enzyme activity of ascorbate peroxidase was determined according to [27] without any modifications. Malondialdehyde content was determined according to [28] with slight modifications. Extract was prepared by the homogenization of 0.1 g of leaf tissue in liquid nitrogen and the addition of 2 ml of 80 % ethanol followed by centrifugation at 14,000 × g at 4 °C for 20 min. In contrast to the original paper, a 0.4 ml aliquot was used for the preparation of the reaction mixture. After heating followed by cooling, the mixture was centrifuged at 1,000 g at 4°C for 20 min.

### 2.4 Hemolytic and cytotoxicity studies

Hemolytic activity was determined according to a previous paper [29]. 1 µl of plant methanol extract (100 mg.ml^−1^) was used per spot. Triton X-100 (1 %, 1 µl per spot, Sigma Aldrich) was used as a known hemolytic agent. The toxicity in mammalian tissue culture was studied on HDF – human dermal fibroblasts, Sigma-Aldrich, 106-05N).

HDF cells were cultivated in DMEM (Sigma-Aldrich) enriched with 10 % fetal bovine serum (Sigma-Aldrich). The cells were maintained in media without antibiotics, however for experiments media supplemented with Antibiotic Antimycotic Solution were used (commercial mixture of penicillin, amphotericin and streptomycin, Sigma-Aldrich).

Cells were harvested from exponential-phase cultures by a standardized detachment procedure using 0.25% Trypsin-EDTA, and the cell number was counted automatically using a Roche’s CASY Cell Counter and Analyzer. 100 ml of 10^5^ cells.ml^−1^ was seeded into the wells for cytotoxicity experiments. Each concentration was tested in quadruplicate within the same experiment in the concentration range 62.5 1,000 µg.ml^−1^. Viability was evaluated after 72 h by standard resazurin assay [30] using fluorescent measurement (560/590nm). Viability was calculated as (sample fluorescence – fluorescence of resazurin) / (fluorescence of cells without treatment - fluorescence of resazurin).

Genotoxicity assay was determined using Hek 293T cells (Human embryonal kidney cells, Sigma-Aldrich, USA) according to Comet assay as previously described [31]. The cells were cultivated as described for HDF cells. For the experiment, 10^5^ cells.ml^−1^ were seeded into the 12-well plates. After 24 hours of cultivation in CO_2_ incubator, the medium was changed and the plant extracts were added to the final concentration of 3 mg.ml^−1^. After 24 hours of incubation, the positive control of genotoxicity was realized by addition of H_2_O_2_ (4.2 mg.l^−1^) for 10 min to the cells. As a negative control served cells without any treatment. After the cells harvesting and electrophoresis, fluorescent microscopy (Olympus IX81 equipped with Texas Red filtr) was used for comets visualization and ImageJ software was used for evaluation TailDNA (%).

### 2.5 Detection of BYDV infection

RNA was isolated by the traditional method using TRIzol reagent (Thermo Fisher Scientific, USA). RT-qPCR assay was performed in a Roche LightCycler® 480 Instrument II using LightCycler® 480 SYBR Green I Master (Roche Applied Science, Germany), RT Enzyme Mix (ArrayScript™ UP Reverse Transcriptase and RNase Inhibitor, Thermo Fisher Scientific) and PVinterF and YanRA primers [32]. The RT-qPCR was performed at 48°C for 30 min, 95°C for 10 min, followed by 40 cycles at 95°C for 15 s and 60°C for 1 min and a melting-curve analysis (95°C 15 s, 60°C 1 min, 95°C 15 s, 60°C 15 s). The qPCR efficiency was determined as E=105.51 %, R2=0.9964 and the linear standard curve interval as 6.89×10^1^ 6.89×10^7^ copies using triplicates for the standard and tested samples. For the standard curve, a specific BYDV nucleotide sequence (294 bp, primers PVinterF+YanRA) amplified by RT-PCR was inserted into the vector pGem-T Easy (Promega) and cloned into *E. coli* JM-109. The selected colony with confirmed insertion sequence was cultivated and DNA extracted (Plasmid Plus Midi Kit, Qiagen, Netherlands) and then excised from the gel (Sigma-Aldrich x-tracta Gel Extraction Tool, Sigma-Aldrich, USA) and purified (GenElute Gel Extraction kit, Sigma-Aldrich). Thereafter, tenfold serial dilution of the transcripts were prepared.

For all samples, the mean detected BYDV concentration was calculated based on the tested triplicates and subsequently normalized using the RNA sample concentration detected spectrophotometrically. For these normalized BYDV titers, their log10 values were calculated and for each week, the log10 mean BYDV titers of the tested group and control group are depicted, together with the interval of plus and minus one standard error of the mean.

### 2.6 Detection of WDV infection

DNA was isolated by adding 0.5 ml of extraction buffer (1 M guanidine thiocyanate, 20 mM Na2H2EDTA, 0.1 M MOPS, pH 4.6, mercaptoethanol added to 0.2% just prior to use) to 50-100 mg of sampled tissue that had been disrupted and homogenized in liquid nitrogen. The solution was incubated for 30 min in a 60°C water bath with occasional vortexing followed by phenol-chloroform-isoamyl alcohol (25:24:1, Affymetrix, USA) extraction, chloroform extraction, isopropanol and sodium acetate precipitation and two steps of 70% ethanol purification. The qPCR assay was performed on a Roche LightCycler® 480 Instrument II using LightCycler® 480 Probes Master (Roche Applied Science, Germany), UniWDVfw and UniWDVrv primers [33] and a TaqMan probe (6-FAM-AGGCGAAGAATGATTCACCCT-BHQ-1). The qPCR was performed at 95°C for 10 min, followed by 40 cycles at 95°C for 15 s and 60°C for 1 min. The qPCR efficiency was determined as E=99.49 %, R^2^=0.9989 and the linear standard curve interval as 2.9×10^3^ – 2.9×10^8^ copies using triplicates for the standard and tested samples. For the standard curve, a specific WDV nucleotide sequence (140 bp, primers UniWDVfw+UniWDVrv) amplified by PCR was inserted into the vector pGem-T Easy (Promega) and cloned into *E. coli* JM-109. The selected colony with confirmed insertion sequence was cultivated and DNA extracted (Plasmid Plus Midi Kit, Qiagen, Netherlands) and then excised from the gel (Sigma-Aldrich x-tracta Gel Extraction Tool, Sigma-Aldrich, USA) and purified (GenElute Gel Extraction kit, Sigma-Aldrich). Thereafter, tenfold serial dilution of the transcripts were prepared.

For all samples, the mean detected WDV concentration was calculated based on the tested triplicates and subsequently normalized using the DNA sample concentration detected spectrophotometrically. For these normalized WDV titers, their log10 values were calculated and for each week, the log10 mean WDV titers of the tested group and control group are depicted, together with the interval of plus and minus one standard error of the mean.

### 2.7 Statistical analysis of experimental data

The statistical significance of results was tested by analysis of variance followed by Duncan’s test in *STATISTICA 12* (data analysis software system, StatSoft. Inc. 2013). For all statistical tests, the significance level was established at p < 0.05.

## 3 Results and discussion

### 3.1 Transgenic barley expressing tobacco osmotin gene

In order to avoid potential mutations originating from the cloning process, the osmotin gene was commercially synthetized *de novo*. The codon optimized osmotin gene was cloned into the expression vector pBract214. The possibility of tobacco osmotin gene expression under the constitutive promoter of the cauliflower mosaic virus has been already by our group demonstrated in barley previously [29]. Therefore, the same promoter was used in this case as well. Transgenic barley plants expressing the osmotin gene were prepared via *Agrobacterium*-mediated transformation. In total, 210 immature embryos were transformed, providing 26 regenerating plants (**Supplementary Figure 2**) of the T0 generation that were transferred into pots and grown to maturity in a greenhouse. The presence of the osmotin transgene was confirmed by PCR in 25 regenerated plants (**Supplementary Figure 3**). The ratio corresponds to a transformation efficiency of 12 %, which is typically lower than for the transformation of dicotyledonous plants [34], but comparable with other barley transformation experiments [35]. Notably, transformation efficiency for *Agrobacterium*-mediated barley immature embryos utilizing hygromycin selection is 25 % in average [20]. Incomplete T-DNA integration into host genomic DNA might occur during *Agrobacterium*-mediated transformation leading to an unintended loss of the selection marker. Previously, it was reported that incomplete integration of T-DNA can reach up to 44 % in monocot wheat [36]. In our experiment, *hpt* selection marker was detected in all transgenic plants thus confirming complete integration of T-DNA cassette (**Supplementary Figure 4**). Heterologous peptide expression can be driven by tissue specific or constitutive promoters. [37] demonstrated usage of root tip specific promoter to induce resistance to biotic stress induced by nematodes. Alternatively, constitutive promoter CaMV35S [38, 39] can be used for heterologous protein production in plants. In our study, expression of osmotin protein was driven by the strong constitutive maize *ubi* promoter that provides strong stable expression in all plant tissues. Broad expression of osmotin should enhance plant response to various biotic and abiotic stresses such as salt stress or fungal infection whose symptoms are not restricted into specific plant tissues but more likely tend to affect the whole plant. The expression of the osmotin gene was confirmed by transgene-specific RT-PCR, while transgene-specific amplicons were not detected in the WT. The specificity of PCR product was additionally verified by melting analysis after performed PCR reaction. The osmotin gene expression was demonstrated in all transgenic plants. There were not observed any visible abnormal phenotypic manifestations of transgenic plants comparing to WT plants suggesting that accumulation of heterologous osmotin protein in plant cells of transgenic barley do not affect substantially plant growth performance. Transgenic plants were prepared for the analysis of the effect of biotic and abiotic stress. First, immature embryos were selected on a medium containing hygromycin. Then, germinating plants were transferred to the pots and the presence of the osmotin transgene was verified by PCR. The segregation ratio of the osmotin transgene in the T1 generation showed Mendelian inheritance. Verified transgenic plants were used in the subsequent experiments.

### 3.2 Higher stress resistance of transgenic barley

Both transgenic and non-transgenic tobacco plants were exposed to stress caused by salinity (200 mM NaCl) and pathogen infection (*Fusarium oxysporum*). As can be seen e.g. in the **Supplementary Figure 5**, the transgenic plants shew a significant reduction in disease symptoms. Both types of stress were able to induce a decrease in protein content in wild-type barley. Many researchers have reported that the level of total soluble protein in crop plants decreases under abiotic stress. Stress usually leads to protein damage caused by *e.g*. reactive oxygen species [40] or by increased activity of proteases [41]. A downregulation of photosynthesis under several stresses was reported at the proteome level [42]. Salt and drought induced a decrease in the main proteins of photosystem II and in both chlorophyll *a* and *b* binding proteins as well as producing a downregulation of RuBisCO and key Calvin cycle enzymes in barley and wheat [43, 44]. However, under stress the transgenic barley expressing the osmotin gene maintained the same protein level as the non-stressed transgenic control plants (**Figure 1**). Similarly, a higher protein level was detected in strawberries recombinantly expressing osmotin [45] in comparison to non-transgenic plants during salt conditions.

**Figure 1.**
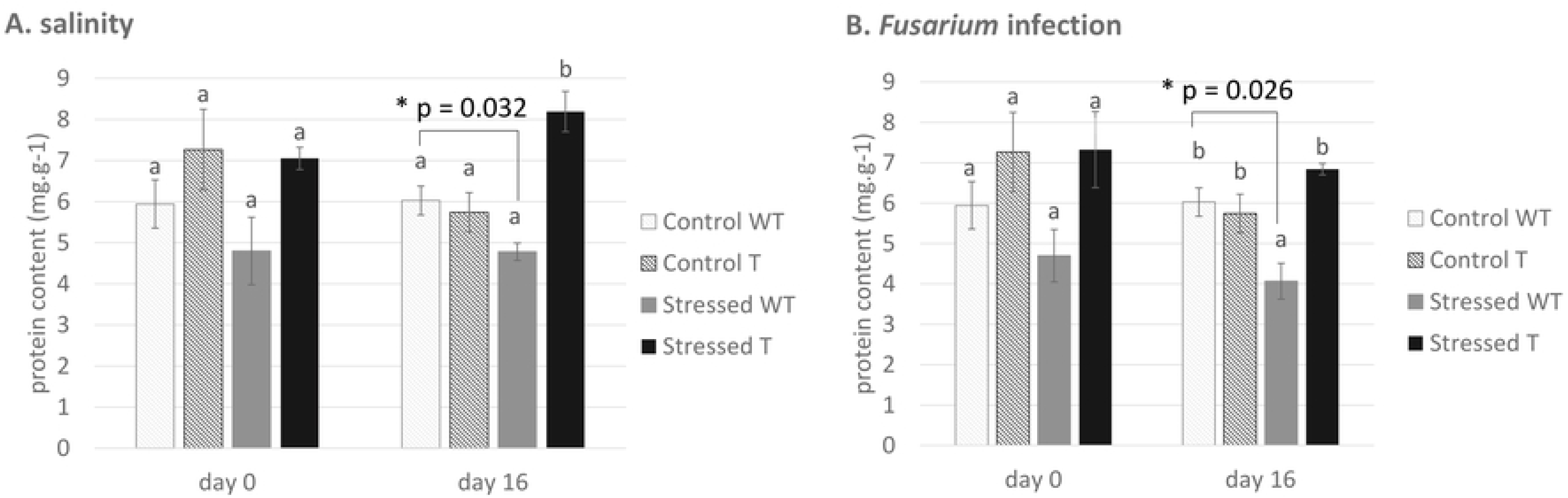
Protein content determined under both types of stress: (a) plants under abiotic stress and (b) plants under biotic stress. WT: non-transgenic (wild type) barley *Hordeum vulgare* L. var. Golden Promise, T: transgenic barley expressing tobacco osmotin gene. The results are shown as the average value of four plants measured in four replicates. Data are presented with the standard error of the mean (SEM). The statistical significance of results was tested by analysis of variance followed by Duncan’s test (p < 0.05). * the groups were compared by two-choice t-test, the p values are presented.

As was mentioned above, photosynthesis is significantly affected during stress conditions; therefore, chlorophyll content in barley was measured in the presence of both types of stressing factors. Ongoing stress was detected in wild-type barley, where both *Fusarium* infection and salinity exhibited an influence on chlorophyll content (**Figure 2**). However, similarly to protein content, the transgenic barley maintained the same chlorophyll level as the control non-stressed plants. In agreement with our results, it has been [46] already reported that transgenic tomato plants expressing the osmotin gene had higher chlorophyll content during the drought and salt stress than the non-transgenic plants. Similarly, osmotin-expressing transgenic soybean, chilli pepper and strawberry exhibited higher chlorophyll content than the non-transgenic variants during salinity [45, 47, 48]. The connection between osmotin and photosynthesis has been already reported, demonstrating an osmotin affinity to brassinosteroids, plant hormones affecting photosynthesis activity [49, 50].

**Figure 2.**
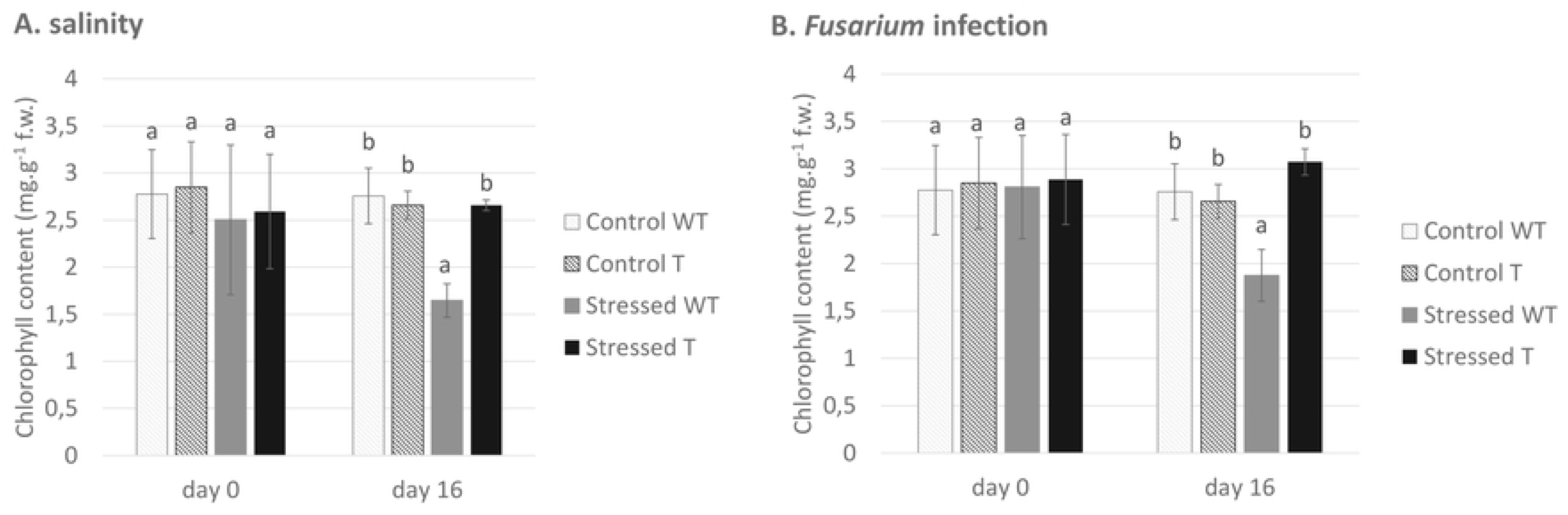
Chlorophyll content determined in both types of stress: (a) plants under abiotic stress and (b) plants under biotic stress. WT: non-transgenic (wild type) barley *Hordeum vulgare* L. var. Golden Promise, T: transgenic barley expressing tobacco osmotin gene. The results are shown as the average value of four plants measured in four replicates. Data are presented with the standard error of the mean (SEM). The statistical significance of results was tested by analysis of variance followed by Duncan’s test (p < 0.05).

As a major part of their defense system, plants have evolved an antioxidant strategy for overcoming stress conditions. Antioxidants (both enzymatic and non-enzymatic) prevent osmotic stress, oxidative stress, molecular damage, and even cell death [51]. Salt stress induces the production of reactive oxygen species (ROS), which causes oxidative stress. Therefore, the amount of antioxidant plays an important role during stressful conditions. Here, attention was focused on APX (ascorbate peroxidase). When influenced by stress, the transgenic barley plants exhibited a higher level of this antioxidant (**Figure 3**), indicating a lower susceptibility to salinity than the non-transgenic control plants. The connection between a higher level of APX and salt stress tolerance was demonstrated in genetically modified sweet potato [52]. Similarly to our results, transgenic chilli pepper and soybean expressing the tobacco osmotin gene exhibited a higher level of APX and improved salt stress tolerance [48, 53]. Lipid peroxidation is a process caused by free radicals (*e.g*. ROS) attacking unsaturated lipids in membranes. Malondialdehyde (MDA) is one of the end products of lipid peroxidation [54]. As we demonstrated that an increased level of APX prevents radical formation in transgenic plants, logically, we also found a lower amount of MDA in transgenic plants during salinity (**Figure 4**). Less MDA, indicating the effect of osmotin on cell membrane protection from damage by lipid peroxidation, has been already reported in transgenic olive plants exposed to drought [55].

**Figure 3.**
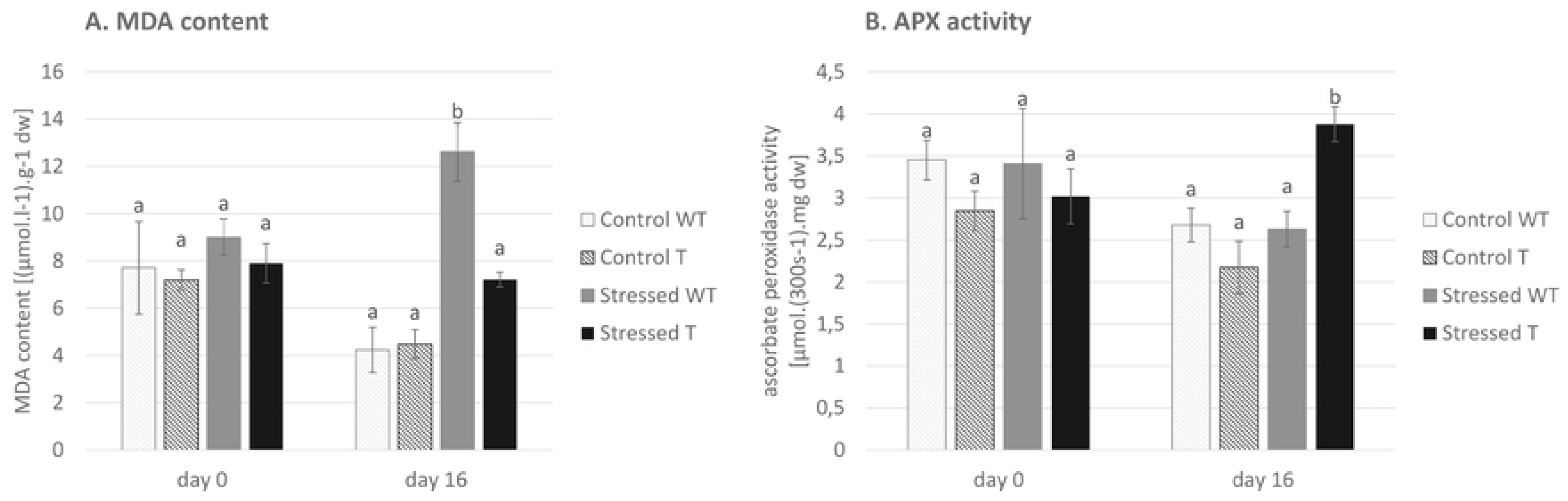
Effect of salinity on lipid peroxidation demonstrated as malondialdehyde (MDA) content (a) and antioxidant activity presented as ascorbate peroxidase (APX) activity (b). WT: non-transgenic (wild type) barley *Hordeum vulgare* L. var. Golden Promise, T: transgenic barley expressing tobacco osmotin gene. The results are shown as the average value of four plants measured in four replicates. Data are presented with the standard error of the mean (SEM). The statistical significance of results was tested by analysis of variance followed by Duncan’s test (p < 0.05).

**Figure 4.**
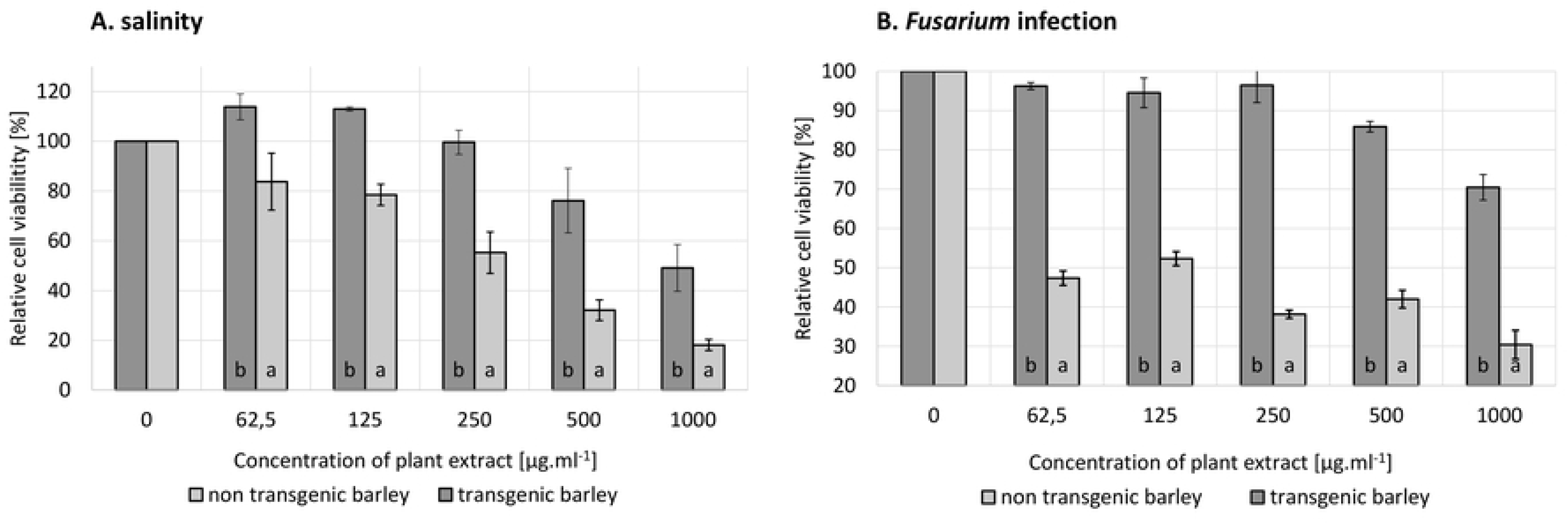
Cytotoxicity of methanol barley extracts on human dermal fibroblasts (HDF). The toxicity was evaluated after 72 hours of cells cultivation over a range of concentrations of extracts by standardized resazurin-based viability assay. The results are shown as the average value of four plants measured in four replicates. Data are presented with the standard error of the mean (SEM). The statistical significance of results was tested by analysis of variance followed by Duncan’s test (p < 0.05).

Both types of stress led to the induction of stress markers such as a decrease in chlorophyll and protein content in wild-type barley plants, however, osmotin-expressing barley plants did not show evidence of ongoing stress, indicating their better preparedness for coping with the stressful conditions. Moreover, during conditions of salt stress, transgenic barley has a higher level of antioxidant and corresponding lower amount of MDA.

### 3.3 Lower toxicity of stressed transgenic barley in comparison to WT

At the end of the exposure to stress, the aboveground biomass was extracted by methanol. The extracts were then added to the growth medium of human fibroblasts in the concentration range from 0 to 1,000 µg.ml^−1^. In both types of stress, there is evidence that transgenic plant extracts are less toxic than those of the non-transgenic (wild-type) ones. The cytotoxicity experiment was done in four technical repetitions for each plant sample, and the viability of fibroblasts (**Figure 5**) was evaluated as the average of four biological repetitions (meaning four independent plants, both transgenic and wild-type variant). The viability of cells decreased with a higher concentration of plant extracts. However, the toxicity of wild-type barley extracts of plants exposed to both types of stress was detected at the lowest tested concentration (62.5 µg.ml^−1^). On the other hand, a toxicity of osmotin-expressing barley extracts was detected at a significantly higher concentration (500 µg.ml^−1^). This finding could confirm our hypothesis that transgenic plants are better prepared for stressful conditions by osmotin expression and therefore do not produce so many secondary metabolites, which are mostly responsible for their toxicity. This finding was confirmed by genotoxicity comparison of transgenic and non transgenic extracts as well where the plants expose to Fusarium

**Figure 5.**
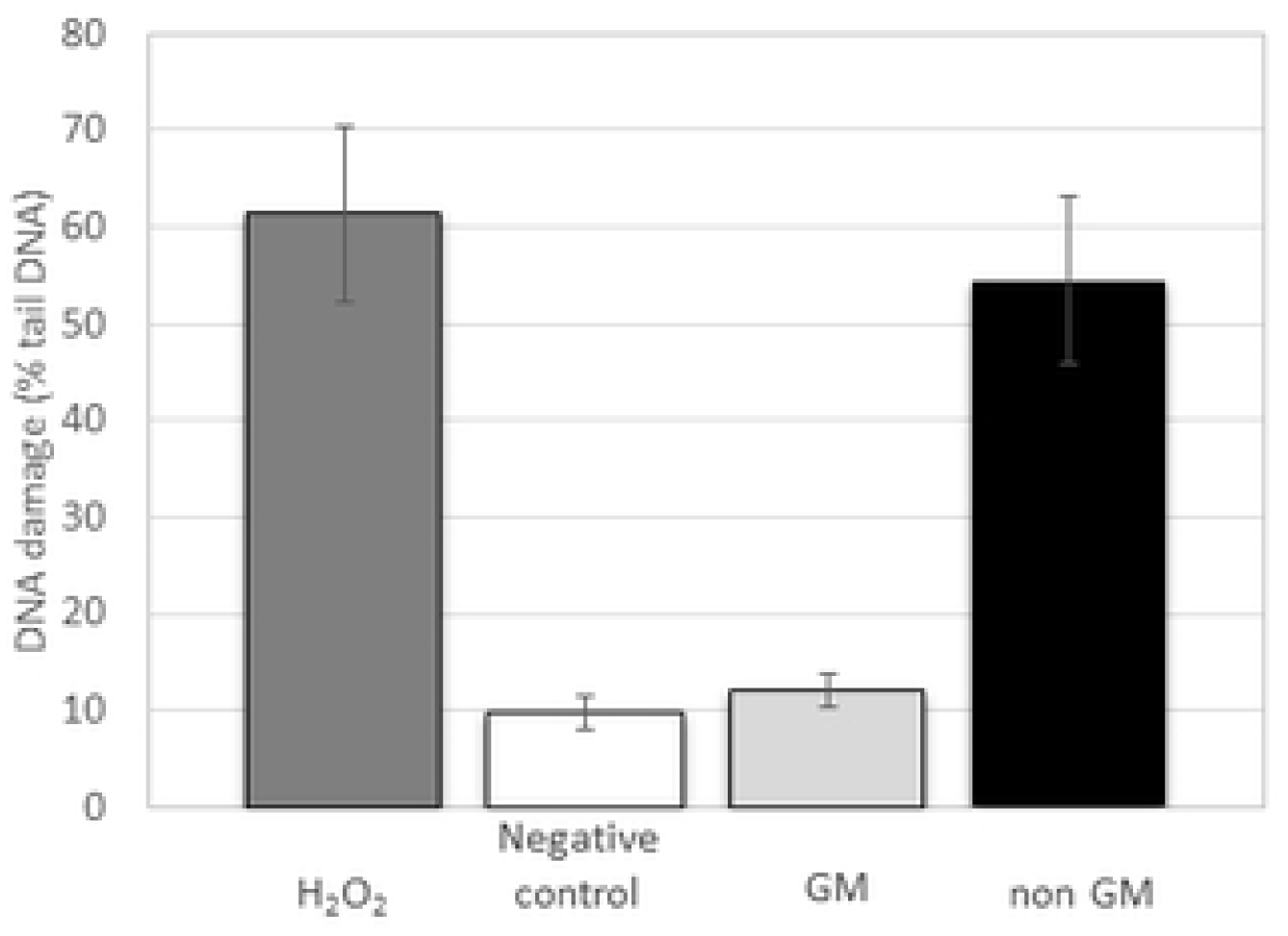
Genotoxicity was evaluated after 24 h of co-cultivation of human embryonic kidney cells with methanol extracts of both transgenic and non transgenic barley exposed to *Fusarium oxysporum* infection. Cells incubated with H_2_O_2_ served as a positive control of DNA damage. Cells without any treatment served as a negative control. Data are presented as the average of 50 individual determinations with the standard error of the mean (SEM).

GM plants, which were modified to cope with environmental stress, have their internal metabolism significantly changed, preventing plant defense system over-response and the accumulation of toxicants, anti-nutrients and secondary metabolites during the ongoing stress [56]. The changes in toxic secondary metabolite content have been already demonstrated by [57] in transgenic potatoes exposed to a pathogen. The genetic manipulation of carbohydrate metabolism and pathogen resistance in these potatoes led to changes in the profile of plant defense compounds, which were mainly characterized by a reduction in the level of the main glycoalkaloids R-solanine and R-chaconine. As well as the expression of plant secondary metabolites, the secondary metabolites formed by the pathogen could have a significant effect on the acute toxicity of crop extracts. In particular, the toxicity of mycotoxins has been reported many times [58]. A lower amount of mycotoxins as a secondary effect of genetic manipulation was detected *e.g*. in a comprehensive study focused on transgenic maize [59]. The mycotoxins, as a secondary metabolite of fungi, could be responsible for the genotoxicity, which we detected in case of methanol extracts from the non transgenic barleys infected by *Fusarium oxysporum*. However, the extracts from transgenic barley expressing the antifungal protein osmotin showed no toxicity in the same test as shown in **Figure 5**. The genotoxicity was evaluated after co-cultivation of plant extracts with human embryonal kidney cells (Hek 293T) by standardized Comet assay with appropriate controls.

### 3.4 Weak or no impact of transgenic barley on viral infection spread by aphids and leafhoppers

The influence of GM crops on biodiversity has been discussed and tested many times (reviewed *e.g*. by [60] and [61]), mostly demonstrating that GM crops have reduced the impacts of agriculture on biodiversity. However, confirmation of this hypothesis is still needed. In this paper, we focused on the effect of barley expressing a multi-functional osmotin protein on virus pathogen – host interactions. For barley the effect of aphids spreading BYDV and the effect of leafhoppers spreading WDV was studied. Both viruses cause worldwide diseases of the most important crops including barley, wheat, rice and maize [62]. As is shown in **Figure 6**, the genetic manipulation of barley by osmotin gene insertion had no effect on obtained virus titres through the whole tested period of first 6 weeks after inoculation. Both aphids and leafhoppers were able to attack GM barley and insert the virus into the phloem and infect the tested plants. For phenotype observations, the difference in plant height and tiller count between wild-type infected and transgenic infected plants was tested 6 weeks after inoculation. For both the WDV and BYDV virus, neither the plant height nor the tiller count after 6 weeks from inoculation was proved to differ. The smallest t-test value for two-sample two-tailed t-test was measured for tiller count for WDV infection (t=0.086), where the infected transgenic barley plants showed smaller, but not significantly different, tiller count (2.29) than the infected wild-type plants (3.38), i.e. exhibiting slightly less severe symptoms of WDV infection, where strongly infected plants often exhibit a high tiller count at the tillering stage. Similarly no impact of potato genetic manipulation on aphids was determined either by [63] or by another research group, which found Bt corn to have no effect on aphids [64].

**Figure 6.**
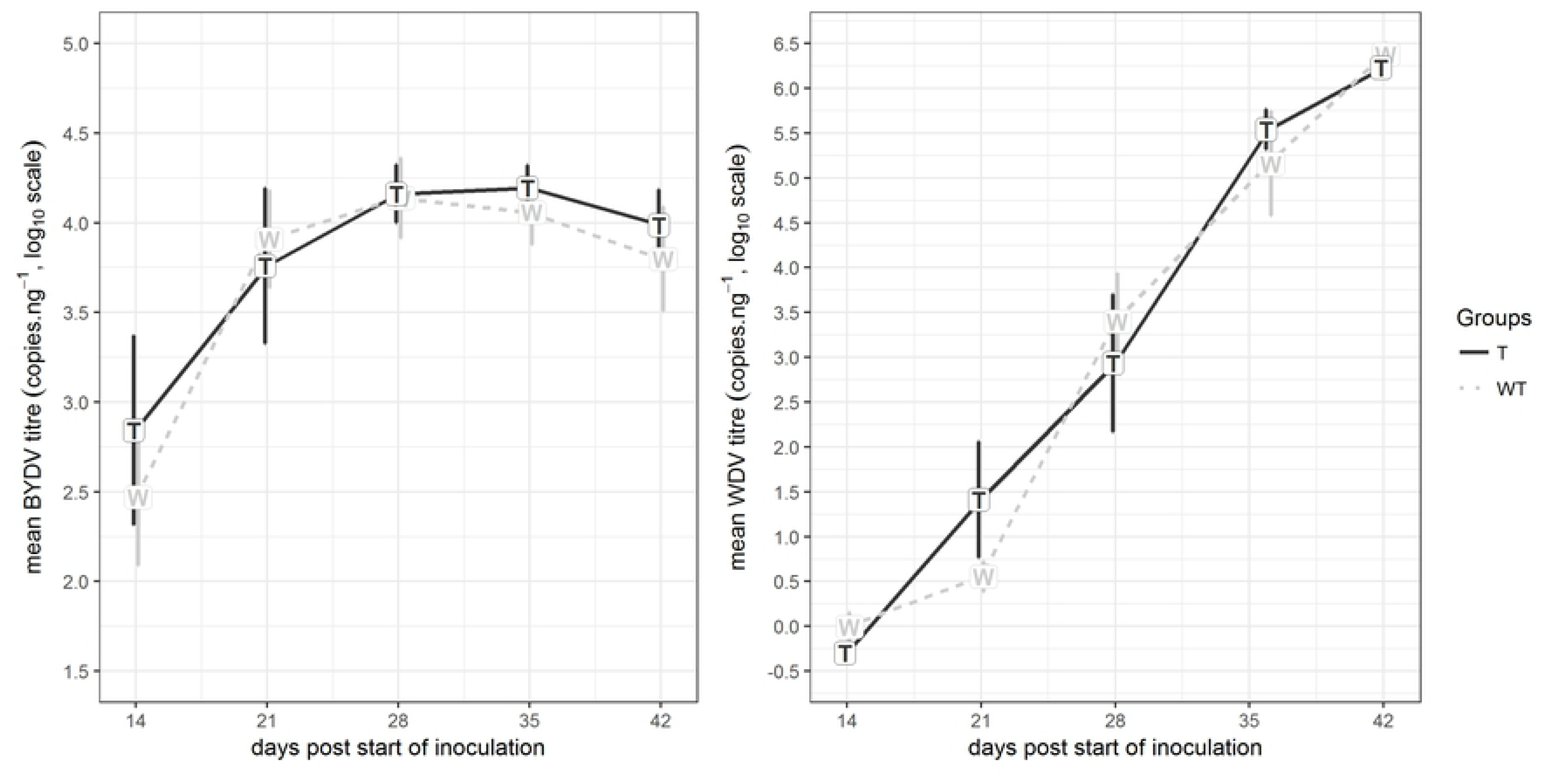
BYDV and WDV titres measured in plant leaves in 42-day period following inoculation (DPI – days post inoculation). BYDV and WDV titres were measured as the mean qPCR identified in titres from triplicates, normalized to (divided by) the RNA sample concentration (for BYDV) or DNA sample concentration (for WDV) and then a base-10 logarithm transformation was applied. From each tested period, the mean from all six tested plants of each group is depicted (WT: non-transgenic (wild type) barley *Hordeum vulgare* L. var. Golden Promise, T: transgenic barley expressing tobacco osmotin gene). Data are presented with the standard error of the mean (SEM). The statistical significance of results was tested by analysis of variance followed by Duncan’s test (p < 0.05).

## 4 Concluding remarks

Many investigations have been carried out to elucidate the mechanisms of the response of the transgenic plant to abiotic and biotic stress, however, the acceptance of transgenic crops in agriculture and industry is still limited, particularly in the EU. We compare osmotin-expressing GM and non-GM barley exposed to biotic and abiotic stress in order to investigate whether their toxicity level under adverse conditions is comparable. The results clearly show that our transgenic barley has a decreased toxicity to human cells under conditions of a/biotic stress for which it is better prepared and exhibits higher stress resistance. These findings provide a new perspective which could help to evaluate the safety of products from genetically modified crops.

## Acknowledgement

*This work was supported by GACR project No. 15-22276S, OPPC CZ.2.16/3.1.00/24503 and NPU I LO1601. The authors thank Ben Watson-Jones, MEng., for providing language corrections. The authors declare no commercial or financial conflicts of interest.*

---

**Supplementary Figure 1** Scheme of the expression vector pBRACT214 containing the osmotin gene.

**Supplementary Figure 2** Regeneration of transgenic barley var. Golden Promise after transformation with osmotin gene. a) Regenerating plantlets from calluses after 6 wk on selection medium. b) Putative transgenic plants on regenerating medium. c) Transgenic plants T0 generation in greenhouse.

**Supplementary Figure 3** Detection of osmotin transgene in the genomic DNA of T0 regenerants. Lane 1-11: samples; lane 12: negative control (DNA/RNA free water); lane 13: positive control (plasmid pBRACT214::osm); lane 14: negative control (genomic DNA of WT plants); lane 15: DNA standard (50 bp DNA ladder, Bioline). Size of PCR product is 222 bp.

**Supplementary Figure 4** Detection of transgene encoding the marker for selection (hpt gene) in the genomic DNA of T0 regenerants. Lane 1-3: samples; lane 4: negative control (DNA/RNA free water); lane 5: negative control (genomic DNA of WT plants); lane 6: positive control (hpt positive plant), lane 7: DNA standard (2-kb DNA ladder, Bioline). Size of PCR product is 275 bp.

**Supplementary Figure 5** left: Non-transgenic barley (left) versus transgenic barley bearing tobacco osmotin gene (right) after biotic stress (15 days after first spraying of Fusarium oxysporum spores). Right: symptoms recognized on non-transgenic barley leaves after stress (chlorosis, necrosis, premature leaf drops and wilt of whole plant).

## Abbreviations

APX: ascorbate peroxidase
BYDV: Barley yellow dwarf virus
GM: genetically modified
HDF: human dermal fibroblasts
MDA: malondialdehyde
ROS: reactive oxygen species
WDV: Wheat dwarf virus
WT: wild type

## References

1. Singh NK, Nelson DE, Kuhn D, Hasegawa PM, Bressan RA. Molecular Cloning of Osmotin and Regulation of Its Expression by ABA and Adaptation to Low Water Potential. Plant physiology. 1989;90(3):1096–101. Epub 1989/07/01. PubMed PMID: 16666857; PubMed Central PMCID: PMCPmc1061849.

2. Singh NK, Handa AK, Hasegawa PM, Bressan RA. Proteins Associated with Adaptation of Cultured Tobacco Cells to NaCl. Plant physiology. 1985;79(1):126–37. PubMed PMID: PMC1074839.

3. Viktorova J, Krasny L, Kamlar M, Novakova M, Mackova M, Macek T. Osmotin, a Pathogenesis-Related Protein. Current Protein & Peptide Science. 2012;13(7):672–81. PubMed PMID: WOS:000311910500008.

4. Liu D, Raghothama KG, Hasegawa PM, Bressan RA. DISEASE RESPONSES TO FUNGAL INFECTION IN TRANSGENIC TOBACCO AND POTATO PLANTS THAT OVEREXPRESS THE OSMOTIN GENE. Plant physiology. 1993;102(1):165-. PubMed PMID: WOS:A1993LD89000953.

5. Parmar N, Singh KH, Sharma D, Singh L, Kumar P, Nanjundan J, et al. Genetic engineering strategies for biotic and abiotic stress tolerance and quality enhancement in horticultural crops: a comprehensive review. 3 Biotech. 2017;7. doi: 10.1007/s13205-017-0870-y. PubMed PMID: WOS:000405242100001.

6. Noori SAS, Sokhansanj A. Wheat plants containing an osmotin gene show enhanced ability to produce roots at high NaCl concentration. Russian Journal of Plant Physiology. 2008;55(2):256–8. doi: 10.1134/s1021443708020143. PubMed PMID: WOS:000254839800014.

7. Rao MVR, Parameswari C, Sripriya R, Veluthambi K. Transgene stacking and marker elimination in transgenic rice by sequential Agrobacterium-mediated co-transformation with the same selectable marker gene. Plant Cell Reports. 2011;30(7):1241–52. doi: 10.1007/s00299-011-1033-y. PubMed PMID: WOS:000291604100009.

8. Beyer P, Al-Babili S, Ye XD, Lucca P, Schaub P, Welsch R, et al. Golden rice: Introducing the beta-carotene biosynthesis pathway into rice endosperm by genetic engineering to defeat vitamin A deficiency. Journal of Nutrition. 2002;132(3):506S–10S. PubMed PMID: WOS:000174189800032.

9. Capellesso AJ, Cazella AA, Schmitt AL, Farley J, Martins DA. Economic and environmental impacts of production intensification in agriculture: comparing transgenic, conventional, and agroecological maize crops. Agroecology and Sustainable Food Systems. 2016;40(3):215–36. doi: 10.1080/21683565.2015.1128508. PubMed PMID: WOS:000374240100003.

10. Talsma EF, Melse-Boonstra A, Brouwer ID. Acceptance and adoption of biofortified crops in low- and middle-income countries: a systematic review. Nutrition Reviews. 2017;75(10):798–829. doi: 10.1093/nutrit/nux037. PubMed PMID: WOS:000412812700003.

11. Qaim M, Kouser S. Genetically Modified Crops and Food Security. Plos One. 2013;8(6). doi: 10.1371/journal.pone.0064879. PubMed PMID: WOS:000320579400029.

12. Nejat N, Mantri N. Plant Immune System: Crosstalk Between Responses to Biotic and Abiotic Stresses the Missing Link in Understanding Plant Defence. Current Issues in Molecular Biology. 2017;23:1–15. doi: 10.21775/cimb.023.001. PubMed PMID: WOS:000405934400001.

13. Al Sinani SSS, Eltayeb EA. The steroidal glycoalkaloids solamargine and solasonine in Solanum plants. South African Journal of Botany. 2017;112:253–69. doi: 10.1016/j.sajb.2017.06.002. PubMed PMID: WOS:000410624200027.

14. Lee ST, Welch KD, Panter KE, Gardner DR, Garrossian M, Chang CWT. Cyclopamine: From Cyclops Lambs to Cancer Treatment. Journal of agricultural and food chemistry. 2014;62(30):7355–62. doi: 10.1021/jf5005622. PubMed PMID: WOS:000339694300005.

15. Breinholt V, Larsen JC. Detection of weak estrogenic flavonoids using a recombinant yeast strain and a modified MCF7 cell proliferation assay. Chemical Research in Toxicology. 1998;11(6):622–9. doi: 10.1021/tx970170y. PubMed PMID: WOS:000074301100005.

16. Di Virgilio AL, Iwami K, Watjen W, Kahl R, Degen GH. Genotoxicity of the isoflavones genistein, daidzein and equol in V79 cells. Toxicology Letters. 2004;151(1):151–62. doi: 10.1016/j.toxlet.2004.04.005. PubMed PMID: WOS:000222226100018.

17. Clifford RJ, Kaplan JH. Human Breast Tumor Cells Are More Resistant to Cardiac Glycoside Toxicity Than Non-Tumorigenic Breast Cells. Plos One. 2013;8(12). doi: 10.1371/journal.pone.0084306. PubMed PMID: WOS:000328734200111.

18. Smith JA, Madden T, Vijjeswarapu M, Newman RA. Inhibition of export of fibroblast growth factor-2 (FGF-2) from the prostate cancer cell lines PC3 and DU145 by Anvirzel and its cardiac glycoside component, oleandrin. Biochemical Pharmacology. 2001;62(4):469–72. doi: 10.1016/s0006-2952(01)00690-6. PubMed PMID: WOS:000169834300010.

19. Alshannaq A, Yu JH. Occurrence, Toxicity, and Analysis of Major Mycotoxins in Food. International Journal of Environmental Research and Public Health. 2017;14(6). doi: 10.3390/ijerph14060632. PubMed PMID: WOS:000404107600080.

20. Harwood WA, Bartlett JG, Alves SC, Perry M, Smedley MA, Leyland N, et al. Barley transformation using Agrobacterium-mediated techniques. Methods in molecular biology (Clifton, NJ). 2009;478:137–47. Epub 2008/11/15. doi: 10.1007/978-1-59745-379-0_9. PubMed PMID: 19009444.

21. Rapacz M, Stępień A, Skorupa K. Internal standards for quantitative RT-PCR studies of gene expression under drought treatment in barley (Hordeum vulgare L.): the effects of developmental stage and leaf age. Acta Physiologiae Plantarum. 2012;34(5):1723–33. doi: 10.1007/s11738-012-0967-1.

22. Kundu JK, Jaro, x, ov, xe, J., et al. Discrimination of Three BYDV Species by One-step RT-PCR-RFLP and Sequence Based Methods in Cereal Plants from the Czech Republic. Cereal Research Communications. 2009;37(4):541–50.

23. J. J, J. C, V. Š, K. KJ. A comparative study of the Barley yellow dwarf virus species PAV and PAS: distribution, accumulation and host resistance. Plant Pathology. 2013;62(2):436–43. doi:doi: 10.1111/j.1365-3059.2012.02644.x.

24. Kundu JK, Gadiou S, Cervena G. Discrimination and genetic diversity of Wheat dwarf virus in the Czech Republic. Virus Genes. 2009;38(3):468–74. Epub 2009/03/28. doi: 10.1007/s11262-009-0352-3. PubMed PMID: 19326201.

25. Tang D, Qian H, Zhao L, Huang D, Tang K. Transgenic tobacco plants expressingBoRS1 gene fromBrassica oleracea var.acephala show enhanced tolerance to water stress. Journal of Biosciences. 2005;30(5):647–55. doi: 10.1007/BF02703565.

26. Bradford MM. A rapid and sensitive method for the quantitation of microgram quantities of protein utilizing the principle of protein-dye binding. Analytical Biochemistry. 1976;72(1):248–54. doi:https://doi.org/10.1016/0003-2697(76)90527-3.

27. Nakano Y, Asada K. Hydrogen Peroxide is Scavenged by Ascorbate-specific Peroxidase in Spinach Chloroplasts. Plant and Cell Physiology. 1981;22(5):867–80. doi: 10.1093/oxfordjournals.pcp.a076232.

28. Hodges DM, DeLong JM, Forney CF, Prange RK. Improving the thiobarbituric acid-reactive-substances assay for estimating lipid peroxidation in plant tissues containing anthocyanin and other interfering compounds. Planta. 1999;207(4):604–11. doi: 10.1007/s004250050524.

29. Viktorova J, Rehorova K, Musilova L, Suman J, Lovecka P, Macek T. New findings in potential applications of tobacco osmotin. Protein Expression and Purification. 2017;129:84–93. doi: 10.1016/j.pep.2016.09.008. PubMed PMID: WOS:000386194500011.

30. Riss TL MR, Niles AL, et al. Assay Guidance Manual. Bethesda (MD): Eli Lilly & Company and the National Center for Advancing Translational Sciences; 2013.

31. Kubásek J, Vojtěch D, Jablonská E, Pospíšilová I, Lipov J, Ruml T. Structure, mechanical characteristics and in vitro degradation, cytotoxicity, genotoxicity and mutagenicity of novel biodegradable Zn–Mg alloys. Materials Science and Engineering: C. 2016;58:24–35. doi:https://doi.org/10.1016/j.msec.2015.08.015.

32. Jarošová J, Kundu JK. Validation of reference genes as internal control for studying viral infections in cereals by quantitative real-time RT-PCR. BMC Plant Biology. 2010;10(1):146. doi: 10.1186/1471-2229-10-146.

33. Gadiou S, Ripl J, Janourova B, Jarosova J, Kundu JK. Real-time PCR assay for the discrimination and quantification of wheat and barley strains of Wheat dwarf virus. Virus genes. 2012;44(2):349–55. Epub 2011/12/17. doi: 10.1007/s11262-011-0699-0. PubMed PMID: 22173982.

34. Sood P, Bhattacharya A, Sood A. Problems and possibilities of monocot transformation. Biologia Plantarum. 2011;55(1):1–15. doi: 10.1007/s10535-011-0001-2. PubMed PMID: WOS:000286669700001.

35. Hisano H, Meints B, Moscou MJ, Cistue L, Echavarri B, Sato K, et al. Selection of transformation-efficient barley genotypes based on TFA (transformation amenability) haplotype and higher resolution mapping of the TFA loci. Plant Cell Reports. 2017;36(4):611–20. doi: 10.1007/s00299-017-2107-2. PubMed PMID: WOS:000398735600009.

36. Wu H, Sparks CA, Jones HD. Characterisation of T-DNA loci and vector backbone sequences in transgenic wheat produced by Agrobacterium-mediated transformation. Molecular Breeding. 2006;18(3):195–208. doi: 10.1007/s11032-006-9027-0.

37. Onyango SO, Roderick H, Tripathi JN, Collins R, Atkinson HJ, Oduor RO, et al. The ZmRCP-1 promoter of maize provides root tip specific expression of transgenes in plantain. Journal of Biological Research-Thessaloniki. 2016;23(1):4. doi: 10.1186/s40709-016-0041-z.

38. Bala M, Thankappan R, Kumar A, Mishra G, Dobraia JR, Kirti PB. Overexpression of a fusion defensin genes from radish and fenugreek improves resistance against leaf spot diseases caused by Cercospora arachidicola and Phaeoisariopsis personata in peanut 2015.

39. Wang Q, Zhu S, Liu Y, Li R, Tan S, Wang S, et al. Overexpression of Jatropha curcas Defensin (JcDef) Enhances Sheath Blight Disease Resistance in Tobacco 2017. 15–21 p.

40. Brady CJ, Gibson TS, Barlow EWR, Speirs J, Wyn Jones RG. Salt-tolerance in plants. I. Ions, compatible organic solutes and the stability of plant ribosomes. Plant, Cell & Environment. 1984;7(8):571–8. doi: 10.1111/1365-3040.ep11591840.

41. Wang X-s, Han J-g. Changes of Proline Content, Activity, and Active Isoforms of Antioxidative Enzymes in Two Alfalfa Cultivars Under Salt Stress. Agricultural Sciences in China. 2009;8(4):431–40. doi:https://doi.org/10.1016/S1671-2927(08)60229-1.

42. Kosová K, Vítámvás P, Prášil IT. Proteomics of stress responses in wheat and barley—search for potential protein markers of stress tolerance. Frontiers in Plant Science. 2014;5:711. doi: 10.3389/fpls.2014.00711. PubMed PMID: PMC4263075.

43. Ghabooli M, Khatabi B, Ahmadi FS, Sepehri M, Mirzaei M, Amirkhani A, et al. Proteomics study reveals the molecular mechanisms underlying water stress tolerance induced by Piriformospora indica in barley. Journal of Proteomics. 2013;94(Supplement C):289–301. doi:https://doi.org/10.1016/j.jprot.2013.09.017.

44. Caruso G, Cavaliere C, Guarino C, Gubbiotti R, Foglia P, Laganà A. Identification of changes in Triticum durum L. leaf proteome in response to salt stress by two-dimensional electrophoresis and MALDI-TOF mass spectrometry. Analytical and Bioanalytical Chemistry. 2008;391(1):381–90. doi: 10.1007/s00216-008-2008-x.

45. Husaini AM, Abdin MZ. Development of transgenic strawberry (Fragaria x ananassa Duch.) plants tolerant to salt stress. Plant Science. 2008;174(4):446–55. doi:https://doi.org/10.1016/j.plantsci.2008.01.007.

46. Goel D, Singh AK, Yadav V, Babbar SB, Bansal KC. Overexpression of osmotin gene confers tolerance to salt and drought stresses in transgenic tomato (Solanum lycopersicum L.). Protoplasma. 2010;245(1-4):133–41. Epub 2010/05/15. doi: 10.1007/s00709-010-0158-0. PubMed PMID: 20467880.

47. Subramanyam K, Sailaja KV, Subramanyam K, Muralidhara Rao D, Lakshmidevi K. Ectopic expression of an osmotin gene leads to enhanced salt tolerance in transgenic chilli pepper (Capsicum annum L.). Plant Cell, Tissue and Organ Culture (PCTOC). 2011;105(2):181–92. doi: 10.1007/s11240-010-9850-1.

48. Subramanyam K, Arun M, Mariashibu TS, Theboral J, Rajesh M, Singh NK, et al. Overexpression of tobacco osmotin (Tbosm) in soybean conferred resistance to salinity stress and fungal infections. Planta. 2012;236(6):1909–25. Epub 2012/09/01. doi: 10.1007/s00425-012-1733-8. PubMed PMID: 22936305.

49. Kamlar M, Uhlík O, Kohout L, Šanda M, Macek T. Affinity chromatography as the method for brassinosteroid-binding protein isolation. J Biotechnol. 2010;150.

50. Rothova O, Hola D, Kocova M, Tumova L, Hnilicka F, Hnilickova H, et al. 24-epibrassinolide and 20-hydroxyecdysone affect photosynthesis differently in maize and spinach. Steroids. 2014;85:44–57. Epub 2014/04/29. doi: 10.1016/j.steroids.2014.04.006. PubMed PMID: 24769061.

51. Caverzan A, Passaia G, Rosa SB, Ribeiro CW, Lazzarotto F, Margis-Pinheiro M. Plant responses to stresses: Role of ascorbate peroxidase in the antioxidant protection. Genetics and Molecular Biology. 2012;35(4 Suppl):1011–9. PubMed PMID: PMC3571416.

52. Yan H, Li Q, Park SC, Wang X, Liu YJ, Zhang YG, et al. Overexpression of CuZnSOD and APX enhance salt stress tolerance in sweet potato. Plant physiology and biochemistry: PPB. 2016;109:20–7. Epub 2016/09/14. doi: 10.1016/j.plaphy.2016.09.003. PubMed PMID: 27620271.

53. Subramanyam K, Sailaja KV, Subramanyam K, Rao DM, Lakshmidevi K. Ectopic expression of an osmotin gene leads to enhanced salt tolerance in transgenic chilli pepper (Capsicum annum L.). Plant Cell Tissue and Organ Culture. 2011;105(2):181–92. doi: 10.1007/s11240-010-9850-1. PubMed PMID: WOS:000289441100005.

54. Saxena R. Chapter 5 - Arthritis as a Disease of Aging and Changes in Antioxidant Status A2 - Preedy, Victor R. Aging. San Diego: Academic Press; 2014. p. 49–59.

55. Silvestri C, Celletti S, Cristofori V, Astolfi S, Ruggiero B, Rugini E. Olive (Olea europaea L.) plants transgenic for tobacco osmotin gene are less sensitive to in vitro-induced drought stress. Acta Physiologiae Plantarum. 2017;39(10):9. doi: 10.1007/s11738-017-2535-1. PubMed PMID: WOS:000411083700017.

56. Safety and nutritional assessment of GM plants and derived food and feed: The role of animal feeding trials. Food and Chemical Toxicology. 2008;46(Supplement 1):S2–S70. doi:https://doi.org/10.1016/j.fct.2008.02.008.

57. Matthews D, Jones H, Gans P, Coates S, Smith LM. Toxic secondary metabolite production in genetically modified potatoes in response to stress. Journal of agricultural and food chemistry. 2005;53(20):7766–76. Epub 2005/09/30. doi: 10.1021/jf050589r. PubMed PMID: 16190629.

58. Bennett JW, Klich M. Mycotoxins. Clinical Microbiology Reviews. 2003;16(3):497–516. doi: 10.1128/CMR.16.3.497-516.2003. PubMed PMID: PMC164220.

59. Ostry V, Ovesna J, Skarkova J, Pouchova V, Ruprich J. A review on comparative data concerning Fusarium mycotoxins in Bt maize and non-Bt isogenic maize. Mycotoxin research. 2010;26(3):141–5. Epub 2010/08/01. doi: 10.1007/s12550-010-0056-5. PubMed PMID: 23605378.

60. Carpenter JE. Impact of GM crops on biodiversity. GM crops. 2011;2(1):7–23. Epub 2011/08/17. doi: 10.4161/gmcr.2.1.15086. PubMed PMID: 21844695.

61. Ammann K. Effects of biotechnology on biodiversity: herbicide-tolerant and insect-resistant GM crops. Trends in Biotechnology. 2005;23(8):388–94. doi:https://doi.org/10.1016/j.tibtech.2005.06.008.

62. Choudhury S, Hu HL, Meinke H, Shabala S, Westmore G, Larkin P, et al. Barley yellow dwarf viruses: infection mechanisms and breeding strategies. Euphytica. 2017;213(8). doi: 10.1007/s10681-017-1955-8. PubMed PMID: WOS:000405440700008.

63. Lazebnik J, Dicke M, ter Braak CJF, van Loon JJA. Biodiversity analyses for risk assessment of genetically modified potato. Agriculture, Ecosystems & Environment. 2017;249(Supplement C):196–205. doi:https://doi.org/10.1016/j.agee.2017.08.017.

64. Lozzia GC, Furlanis C, Manachini B, Rigamonti IE. Effects of Bt corn on Rhopalosiphum padi L. (Rhynchota Aphididae) and on its predator Chrysoperla carnea Stephen (Neuroptera Chrysopidae). 1998.

